# Genetic engineering of hoxb8 immortalized hematopoietic progenitors: a potent tool to study macrophage tissue migration

**DOI:** 10.1101/815043

**Authors:** Solene Accarias, Thibaut Sanchez, Arnaud Labrousse, Myriam Ben-Neji, Aurélien Boyance, Renaud Poincloux, Isabelle Maridonneau-Parini, Véronique Le Cabec

**Affiliations:** Institut de Pharmacologie et Biologie Structurale, IPBS, Université de Toulouse, CNRS, UPS, Toulouse, France

**Keywords:** macrophage migration, conditionally immortalized macrophages, ER-Hoxb8, CRISPR/Cas9

## Abstract

Tumor-associated macrophages (TAM) are detrimental in most cancers. Controlling their recruitment is thus potentially therapeutic. We showed that TAM perform the protease-dependent mesenchymal migration in cancer, while macrophages perform amoeboid migration in other tissues. Inhibition of mesenchymal migration correlates with decreased TAM infiltration and tumor growth, providing rationale for a new cancer immunotherapy specifically targeting TAM motility. To identify new effectors of mesenchymal migration, we produced ER-Hoxb8-immortalized hematopoietic progenitors with unlimited proliferative ability. The functionality of macrophages differentiated from ER-Hoxb8 progenitors was compared to bone marrow-derived macrophages (BMDM). They polarized into M1- and M2-orientated macrophages, generated ROS, ingested particles, formed podosomes, degraded the extracellular matrix, adopted amoeboid and mesenchymal migration in 3D, and infiltrated tumor explants *ex vivo* using mesenchymal migration. We also used the CRISPR/Cas9 system to disrupt gene expression of a known effector of mesenchymal migration, WASP, to provide a proof of concept. We observed impaired podosome formation and mesenchymal migration capacity, thus recapitulating the phenotype of BMDM isolated from *Wasp*-KO mice. Thus, we validate the use of Hoxb8-macrophages as a potent tool to investigate macrophage functionalities.

**Summary statement:** We validate the use of ER-Hoxb8-immortalized hematopoietic progenitors combined to CRISPR/Cas9 technology as a potent tool to investigate macrophage functionalities with a large scale of applications.

## INTRODUCTION

Macrophages are present in all tissues of the organism where they play a central role in the clearance of microorganisms, tissue homeostasis, the mediation of immune and inflammatory responses, and tissue repair. However, tissue infiltration of macrophages also exacerbates pathological processes, such as chronic inflammation, neurodegenerative disorders and cancers (Allavena et al., 2008; Condeelis and Pollard, 2006; Friedl and Weigelin, 2008; Yiangou et al., 2006). Tumor-associated macrophages (TAMs) mainly originate from blood monocytes (Lahmar et al., 2016) and are recruited to the tumor stroma at all stages of cancer progression (Allavena et al., 2008; Condeelis and Pollard, 2006). TAMs can represent more than fifty percent of the tumor mass, and their number positively correlates with poor prognoses in most cancers (Noy and Pollard, 2014; Ruffell and Coussens, 2015). TAMs are involved in several cancer-promoting events such as angiogenesis, lymphangiogenesis, immunosuppression, resistance to therapy and metastasis formation (Noy and Pollard, 2014; Ruffell and Coussens, 2015). Therefore, the control of human macrophage infiltration in cancers is a current therapeutic challenge (Mantovani et al., 2017; Morrison, 2016; Ostuni et al., 2015). However, many questions regarding the mechanisms of macrophage migration remain unresolved and hinders development of novel therapeutic strategies.

Cell migration in tissues occurs in three dimensions (3D), which deeply differs from 2D migration processes (Doyle et al., 2009; Hooper et al., 2006). Human macrophages were shown to share the capacity with only few cell types to use two migration modes in 3D environments: amoeboid and mesenchymal (Van Goethem et al., 2010). The amoeboid movement is characterized by rounded, ellipsoid, or moderately elongated cells that form blebs or generate small actin-rich filopodia (Friedl and Wolf, 2003; Lammermann and Sixt, 2009; Sahai and Marshall, 2003). These cells do not require adhesion to the extracellular matrix (ECM), but use rather a propulsive and pushing migration mode (Fackler and Grosse, 2008; Lammermann et al., 2009; Paluch et al., 2016). This rapid and non-directional motility involves acto-myosin contractions and depends on the Rho–ROCK pathway. Mesenchymal migration consists of cells which present an elongated and protrusive morphology (Fackler and Grosse, 2008; Friedl and Wolf, 2003; Sahai and Marshall, 2003). The movement is slow and directional, involves cell adhesion to the substratum, and requires proteases to degrade the ECM in order to create paths through dense environments. In both human and mouse macrophages, this migration is dependent on podosomes and is not inhibited but rather stimulated by treatment with ROCK inhibitors (Gui et al., 2018; Gui et al., 2014). It has been shown *in vitro* that, in contrast to lymphocytes, neutrophils and monocytes (Cougoule et al., 2012; Friedl and Weigelin, 2008; Lammermann and Germain, 2014), tumor cells or immature human dendritic cells (DCs) can perform these two mechanistically distinct migration modes (Cougoule et al., 2018; Friedl and Wolf, 2003; Sahai and Marshall, 2003). Human and mouse macrophages can also adopt these two migration modes depending on the ECM architecture (Barros-Becker et al., 2017; Cermak et al., 2018; Cougoule et al., 2010; Cougoule et al., 2012; Gui et al., 2018; Gui et al., 2014; Guiet et al., 2012; Jevnikar et al., 2012; Maridonneau-Parini, 2014; Van Goethem et al., 2011; Van Goethem et al., 2010; Verollet et al., 2011; Verollet et al., 2015). *In vivo* in mouse tumors and *ex vivo* in human breast cancer explants, TAM use the protease-dependent mesenchymal migration mode. At the tumor periphery or in inflamed ear derma in mice, macrophages use the amoeboid motility (Gui et al., 2018). Inhibition of matrix metalloproteases (MMPs) blocked the mesenchymal migration of human macrophages ex vivo and mice macrophages in vivo, which correlates with a decrease in both macrophage infiltration into tumors and tumor growth *in vivo*. This provides the rationale for a new strategy in cancer immunotherapy to specifically target TAM through their motility. The use of the first generation of broad spectrum MMP inhibitors in clinics has been tested but proved in the past to be toxic (Overall and Kleifeld, 2006). Thus, identification of specific effectors is a key step in the investigation of new potential therapeutic targets affecting specifically the mesenchymal mode and thus inhibit a migration mode that is used by TAM (and possibly by only a few number of other cells)..

The comprehensive understanding of the mechanisms used by macrophages to migrate through the mesenchymal mode requires an exhaustive approach that could lead to the identification of a large number of potential targets as recently described (Cermak et al., 2018). All these potential effectors now need to be validated as effective actors of macrophage migration both *in vitro* and *in vivo* through functional studies. For such screening approaches, many studies use bone marrow-derived macrophages (BMDMs) from wild-type (WT) and knock-out (KO) mice that have several drawbacks such as the use of numerous animals, the limited number of cells and the impossibility to generate stable mutants in primary cells. Macrophage cell lines such as murine Raw 264.7 cells or human U937, HL-60 or THP1 cells are also widely used, but they are distantly related to blood- or BMDM particularly because they are cancer cells. We therefore intended to establish immortalized macrophages that counters these drawbacks.

Expansion of murine hematopoietic precursors that were transiently immortalized with a retrovirus-delivered and estrogen-inducible form of the transcription factor Hoxb8 have been described (Wang et al., 2006) and validated for the study of hematopoietic cell biology (Cabron et al., 2018; Chu et al., 2019; Di Ceglie et al., 2017; Gurzeler et al., 2013; Hammerschmidt et al., 2018; Lee et al., 2017; Rosas et al., 2011; Wang et al., 2006; Witschi et al., 2010; Zach et al., 2015). The recent coupling of this long-term hematopoietic progenitor cell lines to the CRISPR/Cas9 technology (Hammerschmidt et al., 2018; Roberts et al., 2019) has enabled the creation of new genetically modifiable cell models. During the last few years, expression of Hoxb8 has been used in several studies mainly focused on DC biology (Cabron et al., 2018; Grajkowska et al., 2017; Hammerschmidt et al., 2018; Leithner et al., 2018; Rosas et al., 2011), but only rare studies explored its use to generate surrogate macrophages (Cabron et al., 2018; Roberts et al., 2019; Wang et al., 2006). In addition, only one of these studies directly compared macrophages derived from Hoxb8 progenitors to macrophages derived from bone marrow, and the association of Hoxb8 conditional-immortalization and CRISPR/Cas9 to specifically study macrophage biology in infectious settings (Roberts et al., 2019).

We herein combined the unlimited proliferative capacity of conditionally ER-Hoxb8-immortalized hematopoietic progenitor cells with the CRISPR/Cas9 technology to create a potent tool to identify and investigate the role of new effectors of macrophage functions and particularly their motility. We established ER-Hoxb8 stable progenitors and differentiated them into macrophages. They exhibited the main macrophage functions as compared to BMDMs. As a proof of concept, ER-Hoxb8 stable progenitors were used to obtain macrophages depleted for WASP, an Arp2/3 activator known to be an effector of macrophage motility (Linder et al., 1999). WASP-deficient ER-Hoxb8 cells showed altered mesenchymal migration *in vitro* and *ex vivo* as described in BMDMs of knock-out mice (Park et al., 2014). Therefore, we present here a powerful tool to genetically manipulate macrophages and explore their functionalities in a broad range of applications.

## RESULTS

### Generation of macrophages differentiated from ER-Hoxb8-conditionally-immortalized hematopoietic progenitors and characterization of cell surface differentiation markers, polarization capacity, phagocytosis and the production of reactive oxygen species

ER-Hoxb8-immortalized hematopoietic progenitor cells were generated as described (Wang et al., 2006). Briefly, hematopoietic stem cells were isolated from the bone marrow of WT C57BL/6 mice and infected with an Estrogen Receptor(ER)-regulated Hoxb8 retrovirus allowing proliferation of macrophage progenitors in the presence of estrogen. These immortalized progenitors had a high proliferation rate (Fig S1A) and could be stably maintained in culture for four months. After selection of stably transduced progenitors, differentiation into macrophages was induced by estrogen removal and addition of M-CSF. To characterize the capacity of these immortalized progenitors to differentiate into fully functional macrophages (called Hoxb8-macrophages thereafter), we compared their characteristics to BMDMs in parallel experiments.

First, cell morphology was observed by scanning electron microscopy. As shown in Fig. 1A, while Hoxb8 progenitor cells showed a round morphology and poorly adhered to the substrate, Hoxb8-macrophages were flat and adherent like BMDMs. Spreading of Hoxb8-macrophages and BMDM on coverslips were comparable (Fig. 1B).

**Figure 1:**
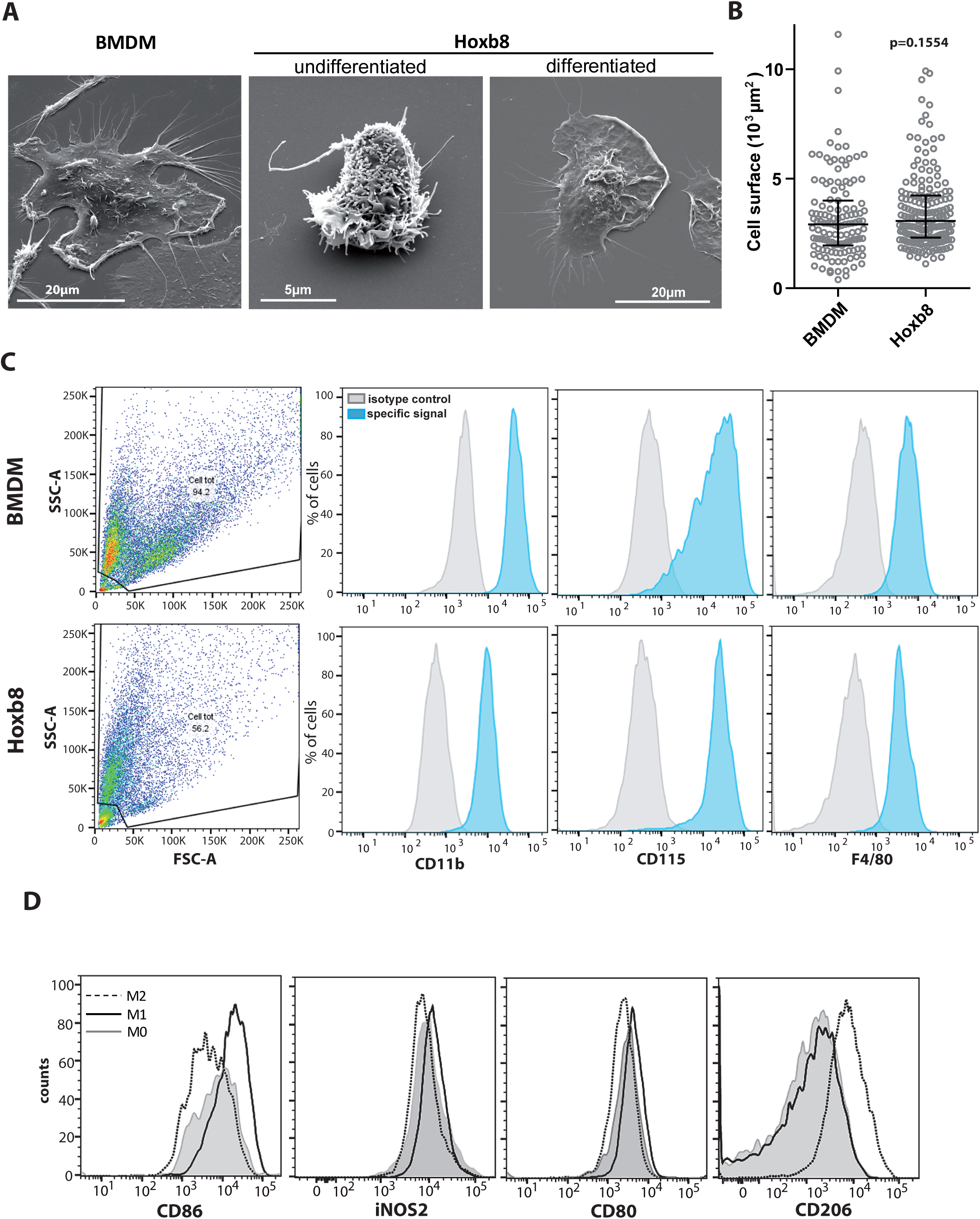
BMDMs and Hoxb8-macrophages share the same morphology, molecular markers and polarization capacity. **(A)** Undifferentiated or macrophage-differentiated ER-Hoxb8 cells and BMDMs were cultured on glass coverslips for 16 h and fixed before being imaged by SEM. Images representative of three independent experiments. **(B)** Cell surface of BMDMs and Hoxb8 macrophages cultured on glass coverslips were measured using the Fiji software on at least 100 cells from 4 independent coverslips. Scatter plot with median and interquartile range is shown. Unpaired student *t*-test p-value is indicated. **(C)** Hoxb8-macrophages and BMDM were analyzed by FACS for their surface expression of 3 specific macrophage markers: CD11b, CD115 and F4/80. Left panels: representative scatter plots of SSC vs FSC. Indicated antibody specific labeling (blue) compared to background (grey). Results are representative of four experiments. **(D)** Expression of CD86, iNOS2, CD80 and CD206 was analyzed by FACS in M0- (grey line), M1- (black line) and M2- (dashed line) polarized Hoxb8-macrophages. Histograms representative of three experiments.

Next, differentiation of Hoxb8 cells was investigated by analysis of surface expression of macrophage-specific markers by FACS. As shown in Fig. 1C, both Hoxb8-macrophages and BMDMs expressed CD11b, CD115 and F4/80 at comparable levels.

To test the capacity of Hoxb8-macrophages cells to polarize into pro- (M1) or anti- (M2) inflammatory macrophages, we analyzed their morphology and surface expression of standard M1/M2 markers described for macrophages (Mantovani et al., 2013; McWhorter et al., 2013; Sica and Mantovani, 2012). When incubated overnight with LPS and γIFN to induce M1 polarization, Hoxb8-macrophages spread (Supplemental Figure S1B) and expressed higher levels of CD86, iNOS and CD80 than M0 or M2 cells (Fig. 1D). When incubated with IL-4 to induce M2 polarization, they acquired an elongated morphology (Supplemental Figure S1B) and the expression of CD206 was enhanced (Fig. 1D).

Next, the microbicidal functions of Hoxb8-macrophages were examined. Reactive oxygen species (ROS) production was measured in response to a phagocytic process triggered by Zymosan particles. ROS production was assessed by fluorescence microscopy using a cell-permeant reactive oxygen species (ROS) detection agent (Fig. 2A). As shown on Fig 2A, while ROS production was poorly detected in resting Hoxb8-macrophages and BMDMs, it was enhanced in both cell types after stimulation with Zymosan. Of note, the signal was enriched around phagosomes (Fig. 2A zoom arrow), as previously described (DeLeo et al., 1999). Similar results were obtained with the cytochrome C reduction assay used to detect superoxide anion production (Le Cabec and Maridonneau-Parini, 1995) (Fig. 2B). Phagocytosis of FITC-Zymosan was next measured by fluorescence microscopy (Fig. 2C, D). Again, the phagocytic capacity of Hoxb8-macrophages was comparable to that of BMDMs with similar percentages of phagocytic cells and similar number of ingested particles per cell (Fig 2C)., as recently described for Mycobacterium tuberculosis and Listeria monocytogenes (Roberts et al., 2019).

**Figure 2:**
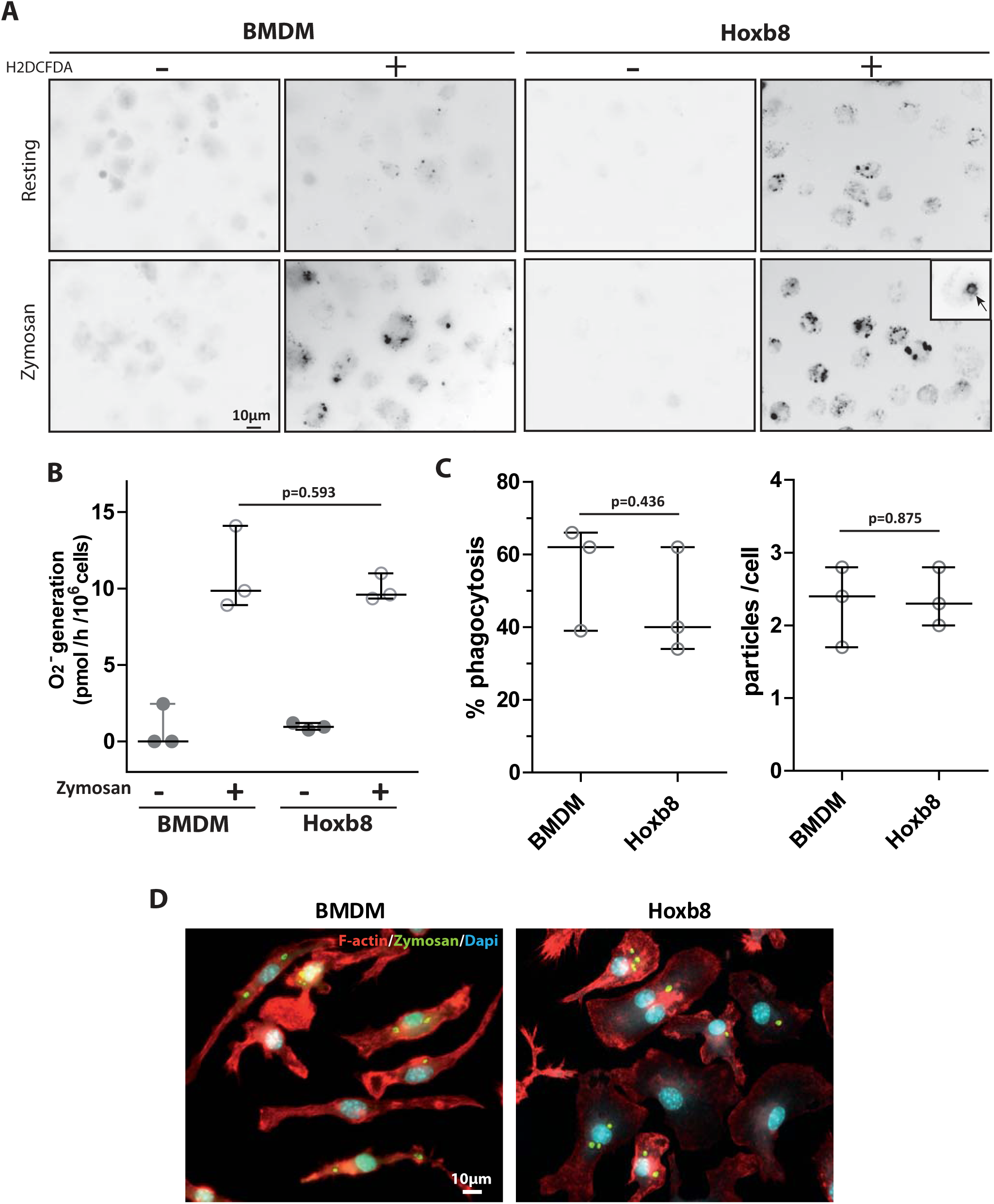
BMDMs and Hoxb8-macrophages have comparable NADPH oxidase and phagocytic activities. BMDMs and Hoxb8-macrophages were cultured on glass coverslips over-night. **(A)** Intracellular ROS generation was measured in parallel in resting or zymosan-activated cells in the presence or absence of H2DCFDA. Fluorescent signals were acquired using a Leica upright DMLB microscope. Images representative of three experiments. **(B)** Quantification of ROS generation by cytochrome C reduction was performed. Median with interquartile of three experiments. Unpaired student *t*-test p-value is indicated. **(C, D)** BMDMs or Hoxb8-macrophages were incubated for 5 h with FITC-Zymosan particles (MOI of 2 particles per cell). Cells stained with rhodamine-phalloidine and DAPI are shown. (C) The percentage of phagocytosis (n=100 cells) and the number of ingested particles per cell were determined manually using the Fiji software in three independent experiments. Unpaired student *t*-test p-value is indicated. (D) Representative images of both cells types with ingested zymosan.

Together, these results indicate that the immortalized Hoxb8 cells efficiently differentiate into macrophages with comparable polarization, phagocytosis and ROS production capacities to BMDMs.

### Characterization of migration, podosome formation and ECM degradation capacities of Hoxb8-macrophages

Since our objective is to use Hoxb8-macrophages as a tool to determine the molecular mechanisms involved in macrophage tissue infiltration, we next analyzed their migration capacities. Hoxb8-macrophages adhered to the substrate and migrated in 2D similarly to BMDMs (Fig. S1C). In the 3D migration assay, Hoxb8-macrophages migrated inside matrices and adopted the expected elongated and protrusive phenotype of mesenchymal migration in Matrigel and a round morphology characteristic of amoeboid migration in fibrillar collagen similarly to BMDMs (Fig. 3A, and supplemental movie 1 to 3). Hoxb8-macrophage infiltration in fibrillar collagen was comparable to that of BMDMs with 41.7+/−3.1% and 47.9+/−2.3%, respectively (mean+/−SD, n=2 to 4). In Matrigel, it was slightly reduced from 59.3+/−2.3 (mean+/−SD n=4) for BMDMs to 38.2+/−9.3% (mean+/−SD n=3) for Hoxb8-macrophages (Fig. 3B). Migration distances inside matrices were comparable in fibrillar collagen (1110+/−349 µm for Hoxb8-macrophages versus 757+/−413 µm for BMDMs, mean+/−SD n=2 to 4) and slightly reduced in Matrigel (577.5+/−86.2 µm for Hoxb8-macrophages *versus* 890+/−141 µm for BMDMs)(Fig 3B).

**Figure 3:**
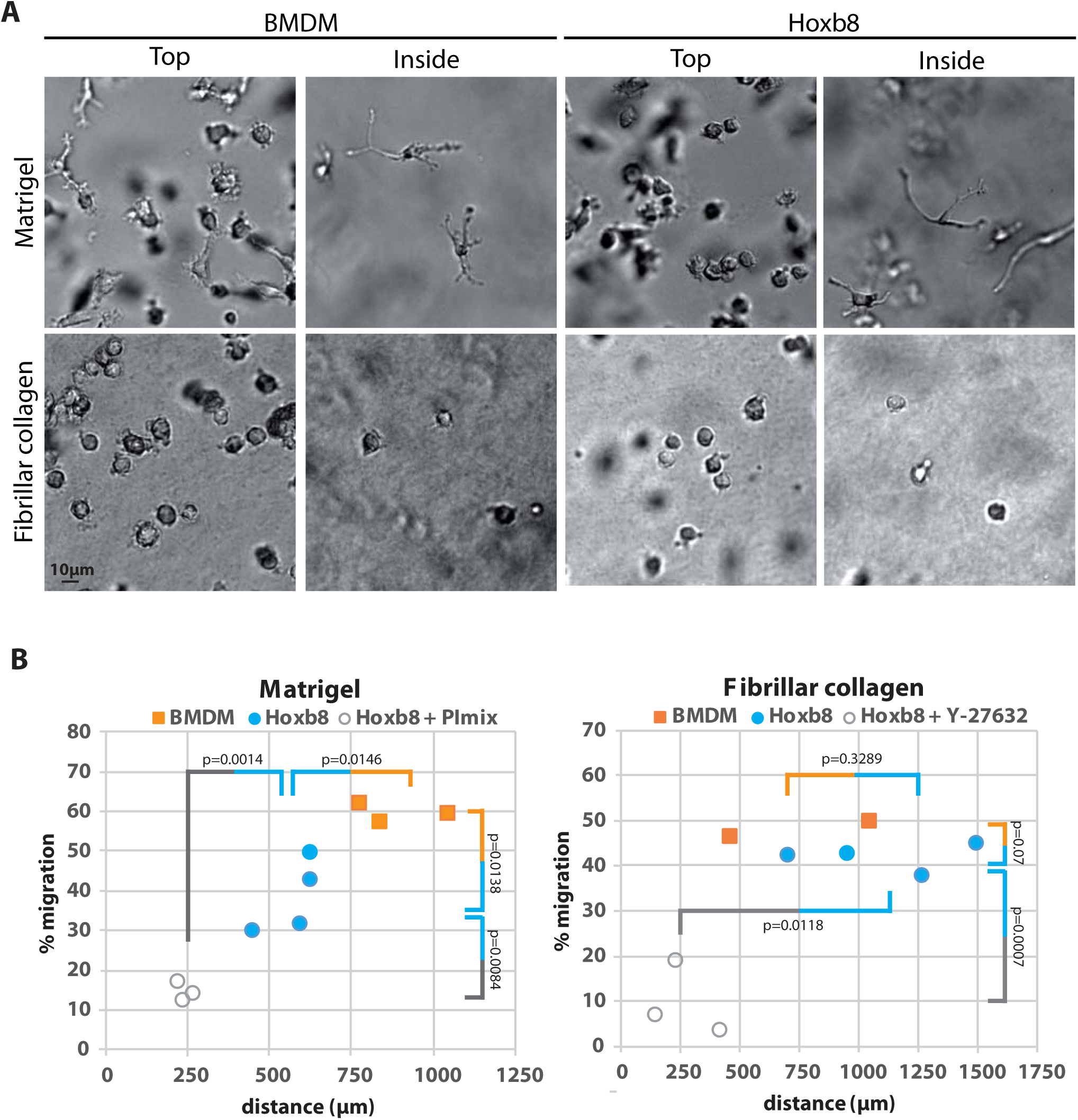
BMDMs and Hoxb8-macrophages have comparable migration capacities in 3D and can perform both amoeboid and mesenchymal migration modes *in vitro*. 3D migration through Matrigel and fibrillar collagen was measured. **(A)** Morphology of Hoxb8-macrophages and BMDMs at the top and inside both extracellular matrices is shown. Representative images of three independent experiments. **(B)** Percentages of cell migration plotted as a function of distance of migration in control conditions for BMDMs (orange square) and Hoxb8-macrophages (blue squares) or in the presence of indicated inhibitors for Hoxb8-macrophages (grey circles)(PI mix in Matrigel or Y-27632 in fibrillar collagen) were measured. Mean of duplicates of 4 independent experiments. Statistics: p-values obtained with paired student *t*-test to compare % of migration (vertical bars) and distances (horizontal bars).

Mice and human macrophage mesenchymal migration in Matrigel depends on protease activity and is therefore inhibited by a mix of proteases inhibitors (PI mix) (Gui et al., 2018; Gui et al., 2014; Guiet et al., 2011; Jevnikar et al., 2012; Van Goethem et al., 2010). In contrast, amoeboid migration in fibrillar collagen is dependent on the Rho/ROCK pathway and is therefore inhibited by the ROCK inhibitor Y-27632 (Gui et al., 2018; Gui et al., 2014; Guiet et al., 2011; Jevnikar et al., 2012; Van Goethem et al., 2010). To ascertain that Hoxb8-macrophages used the mesenchymal migration in Matrigel and the amoeboid migration in fibrillar collagen, as expected for macrophages, PI mix or Y-27632 were used in the migration assay. As shown in Fig. 3B, Hoxb8-macrophages migration in Matrigel and fibrillar collagen were significantly inhibited by the PI mix and Y-27632 respectively.

Therefore, migration of Hoxb8-macrophages matches characteristics of mesenchymal and amoeboid migration when migrating in 3D in Matrigel and fibrillar collagen respectively, with regard to cell morphology, cell dynamic and drug responsiveness (Gui et al., 2018; Van Goethem et al., 2010).

Since the capacity of macrophages to perform mesenchymal migration is correlated to their capacity to form podosomes and to degrade the ECM (Cougoule et al., 2018; Cougoule et al., 2012; Maridonneau-Parini, 2014), we then analyzed the presence of this specialized structures in Hoxb8-macrophages as previously described (Cabron et al., 2018), and compared their organization to BMDMs. Podosomes are submicrometer adhesion cell structures formed at the plasma membrane that protrude into, probe, and degrade the extracellular environment by releasing proteases (Labernadie et al., 2014; Linder and Kopp, 2005; Wiesner et al., 2014). They appear during macrophage differentiation and are not present in monocyte progenitors or in monocytes (Cougoule et al., 2012). They are dynamic cell structures with different spatial organizations (Poincloux et al., 2006; Van Goethem et al., 2011). In resting human macrophages they are mainly scattered (Poincloux et al., 2006; Van Goethem et al., 2011). In BMDMs, they more often assemble as podosome clusters (Fig4A BMDM upper picture) or rosettes (Fig4A BMDM lower picture) as previously described (Cougoule et al., 2010). As shown in Fig 4A, Hoxb8-macrophages exhibited podosome clusters and rosettes in the same proportion as BMDMs. Consequently, they were able to readily degrade the FITC-gelatin, although less efficiently than BMDMs (Fig 4B).

**Figure 4:**
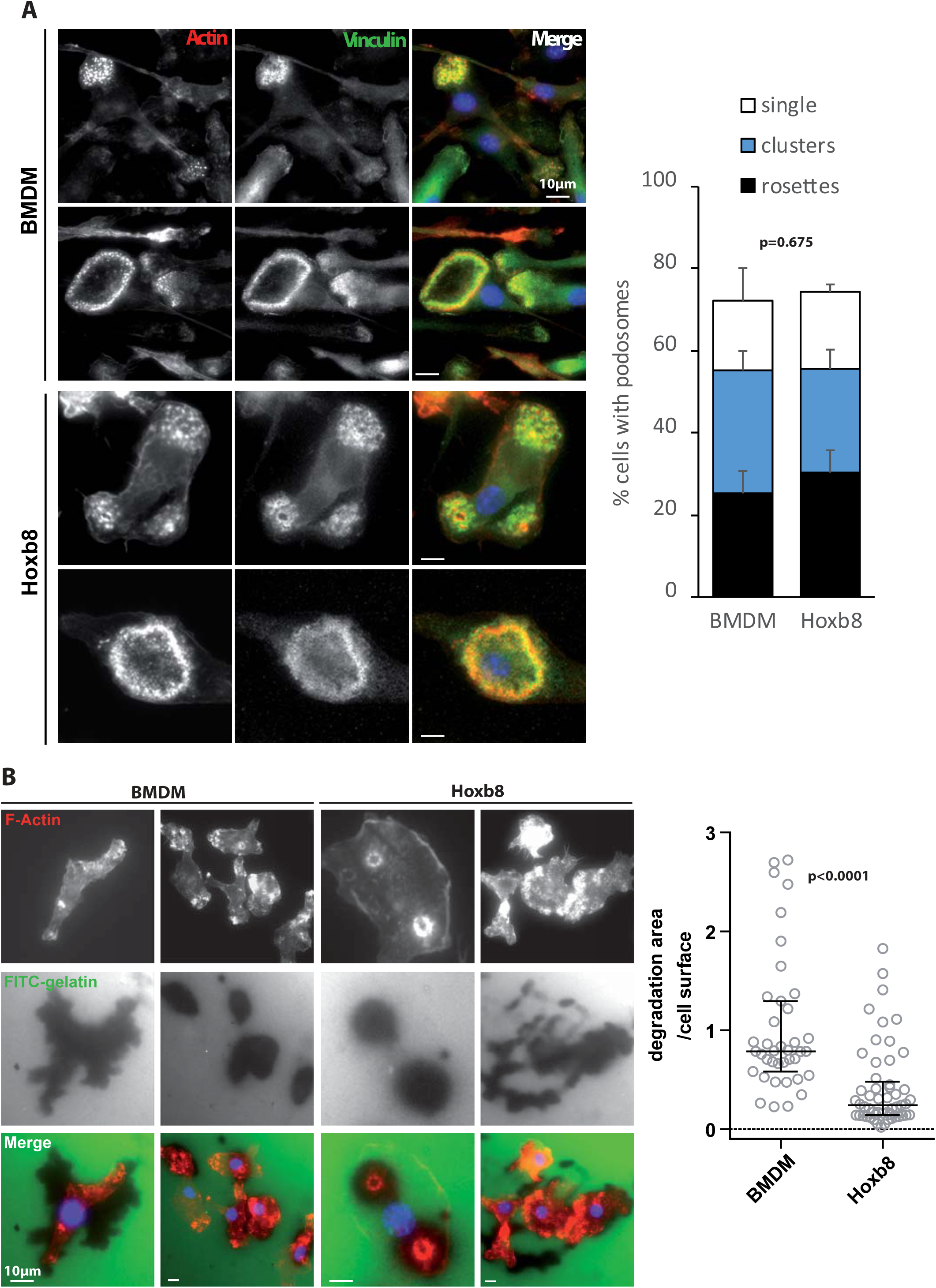
Hoxb8-macrophages and BMDMs have comparable podosomes and ECM degradation capacities. **(A)** Cells were cultured on glass coverslips for 16 h before being processed for the labelling of F-actin, vinculin and the nuclei. Representative pictures of three independent experiments. The percentages of cells with single podosomes, podosome clusters and podosome rosettes in each cell type was quantified on at least 75 cells in 3 independent experiments. Statistics: unpaired student t-test p-value is indicated. **(B)** Cells were seeded on FITC-gelatin coated coverslips. After 24 h, cells were fixed and degradation area were observed by fluorescent microscopy. Quantification of the degradation area was performed using the Fiji Software on 50 cells of 6 independent coverslips. The ratio of degradation area over cell surface were determined and is expressed as median with interquartile range. Statistics: the p-value obtained with an unpaired student *t*-test is shown.

Finally, we tested the capacity of Hoxb8-macrophages to infiltrate tumoral tissue *ex vivo* and *in vivo* (Gui et al., 2018). For *ex vivo* experiments, fibrosarcoma were generated by subcutaneous injection of LPB tumor cells (Gui et al., 2018), tumors were resected and sliced. Cell tracker-labeled BMDMs or Hoxb8-macrophages were layered on tumor slices and co-cultured for three days. The effects of BB-94, a broad-spectrum MMP inhibitor, and Y-27632 were explored in parallel using tissue slices from the same tumor. As shown in Fig 5A, Hoxb8-macrophages were able to infiltrate tumor slices very efficiently as compared to BMDMs. Similarly to BMDMs, their infiltration into tissue was significantly inhibited by BB-94 but not Y-27632 (Fig. 5B) showing that both BMDMs and Hoxb8 macrophages use the mesenchymal mode to infiltrate tumor explants as previously described for human macrophages (Gui et al., 2018). For in vivo experiments, fibrosarcoma were generated in a surgically implanted dorsal window chamber as previously described (Gui et al., 2018). Hoxb8 macrophages pre-labeled with cell tracker were co-injected and their displacement in the tumor was observed by intravital microscopy. As shown in the movie (supplemental movie 4), a mesenchymal displacement is again observed, confirming the results obtained in vitro and ex vivo.

**Figure 5:**
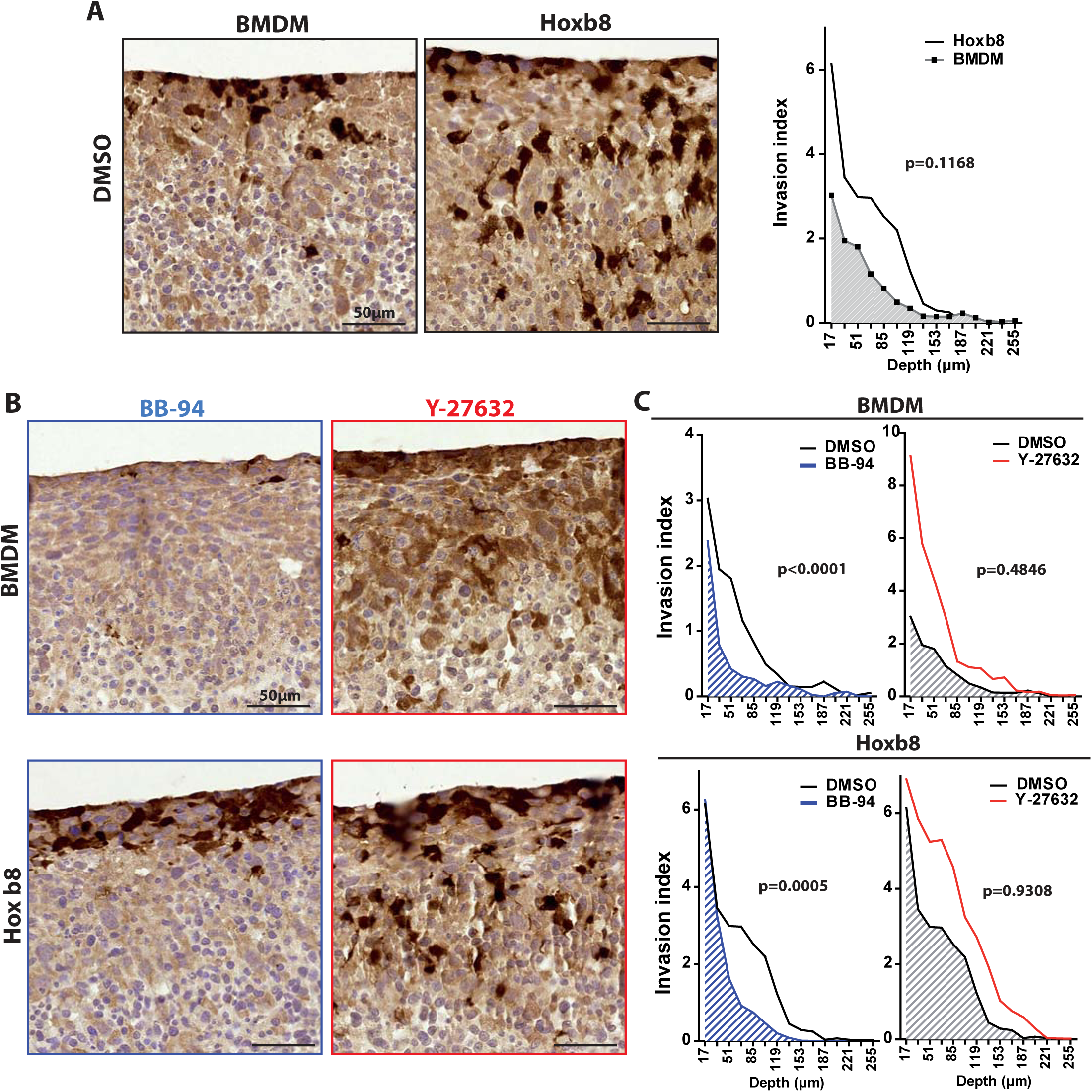
Hoxb8-macrophages and BMDMs infiltrate tumor explants using mesenchymal migration mode *ex vivo*. Cell tracker-labelled cells were seeded on top of sliced LPB tumor explants in the presence of DMSO **(A)** or in the presence of BB-94 (10 µM) or Y-27632 (20 µM) **(B)** over 3 days. Slices were then fixed and serial sectioning was performed along the Z axis. Sections were stained for IHC with an anti-cell tracker antibody to estimate macrophage tissue infiltration and counterstained with hematoxylin. Representative pictures are shown. Scale bar, 50 µm. For all experimental conditions, the invasion index was calculated as the percentage of brown area (cell tracker labelled-cells) over total area and was ploted as a function of invasion depth inside tumor slices. Four tumors were used and means are shown. Statistics: the p-value obtained with an unpaired student *t*-test is shown.

Altogether, these results demonstrate that Hoxb8-macrophages represent a relevant model to study podosome formation, ECM degradation and macrophage migration amongst other functions.

### Generation and characterization of CRISPR/Cas9 WASP-depleted Hoxb8-macrophages

To explore whether the Hoxb8 cell line is a valuable tool to study molecular mechanisms of macrophage tissue migration, we decided to silence the expression of a known effector of macrophage mesenchymal migration, the Wiskott-Aldrich syndrome protein (WASP) (Calle et al., 2008; Linder et al., 1999; Park et al., 2014), using the CRISPR/Cas9 technology. WASP is a hematopoietic cell specific protein and an actin nucleation promoting factor, which regulates Arp2/3-dependent actin polymerization (Takenawa and Suetsugu, 2007). WASP activity is required for several macrophage functions, including podosome formation, chemotaxis and mesenchymal migration but not amoeboid migration (Linder et al., 1999; Park et al., 2014).

WASP-depleted Hoxb8-macrophages (called *Wasp*−/−) were generated using the lentiCRISPRv2 transfer plasmid containing a gRNA sequence targeting exon 1 of the *Wasp* gene. As a control, cells were transduced with “empty” lentiviruses expressing Cas 9 but devoid of gRNA and called *Wasp* +/+.

As shown in Fig 6A, WASP was efficiently depleted by 91.6 +/−2.2% (mean+/−SD, n=3) in *Wasp*−/− Hoxb8 macrophages as compared to *Wasp* +/+ or the parental Hoxb8 macrophages. WASP depletion was not compensated by the up-regulation of N-WASP expression (Supplemental Fig. S1D) as previously described in *Wasp*-knock out (KO) mice (Isaac et al., 2010).

**Figure 6:**
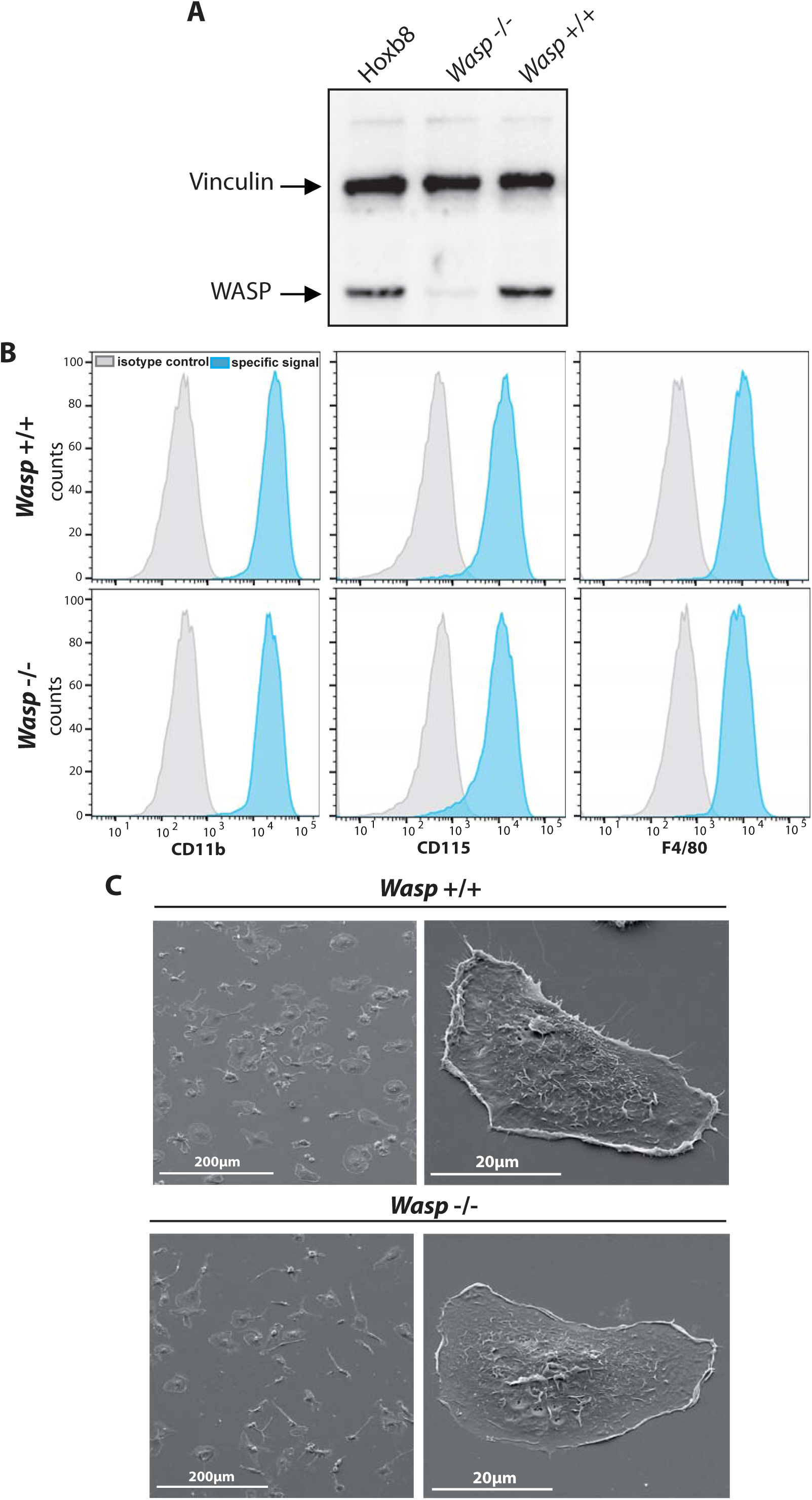
Establishment and characterization of WASP-depleted Hoxb8-macrophages. (A) Expression level of WASP in original Hoxb8-, *Wasp* +/+ and *Wasp* −/− Hoxb8-macrophages was analyzed by western blot. **(B)** Expression of characteristic macrophage markers were analyzed by FACS on *Wasp* +/+ and *Wasp* −/− Hoxb8-macrophages as described in Fig. 1. One representative experiment out of four is shown. **(C)** Morphology of *Wasp* +/+ and *Wasp* −/− was analyzed by SEM as described in Fig. 1. Representative images of three experiments. No obvious differences in cell morphology or adhesion was noticed.

We first checked whether extinction of WASP impacted Hoxb8 cells differentiation. As shown in fig 6B, the expression of CD11b, CD115 and F4/80 was comparable between *Wasp* +/+ and *Wasp*−/− Hoxb8 macrophages. Cell morphology and spreading of *Wasp* +/+ and *Wasp* −/− Hoxb8 macrophages were also similar (Fig. 6C). Podosome rosettes appeared similar in both *Wasp* +/+ and *Wasp* −/− macrophages, but podosome clusters appeared disorganized and less abundant in *Wasp* −/− cells (Fig 7A, B). Also, the global percentage of cells with podosomes and the proportion of single podosomes and podosome clusters were significantly reduced in *Wasp*−/− Hoxb8 macrophages compared to control cells (Fig 7B). The ability of *Wasp* −/− Hoxb8 macrophages to degrade FITC-gelatin was significantly reduced (Fig. 7C, D), as was their capacity to migrate using the mesenchymal mode (Fig 8A), thus recapitulating results obtained with BMDMs isolated from *Wasp* knock out mice (Calle et al., 2004; Park et al., 2014).

**Figure 7:**
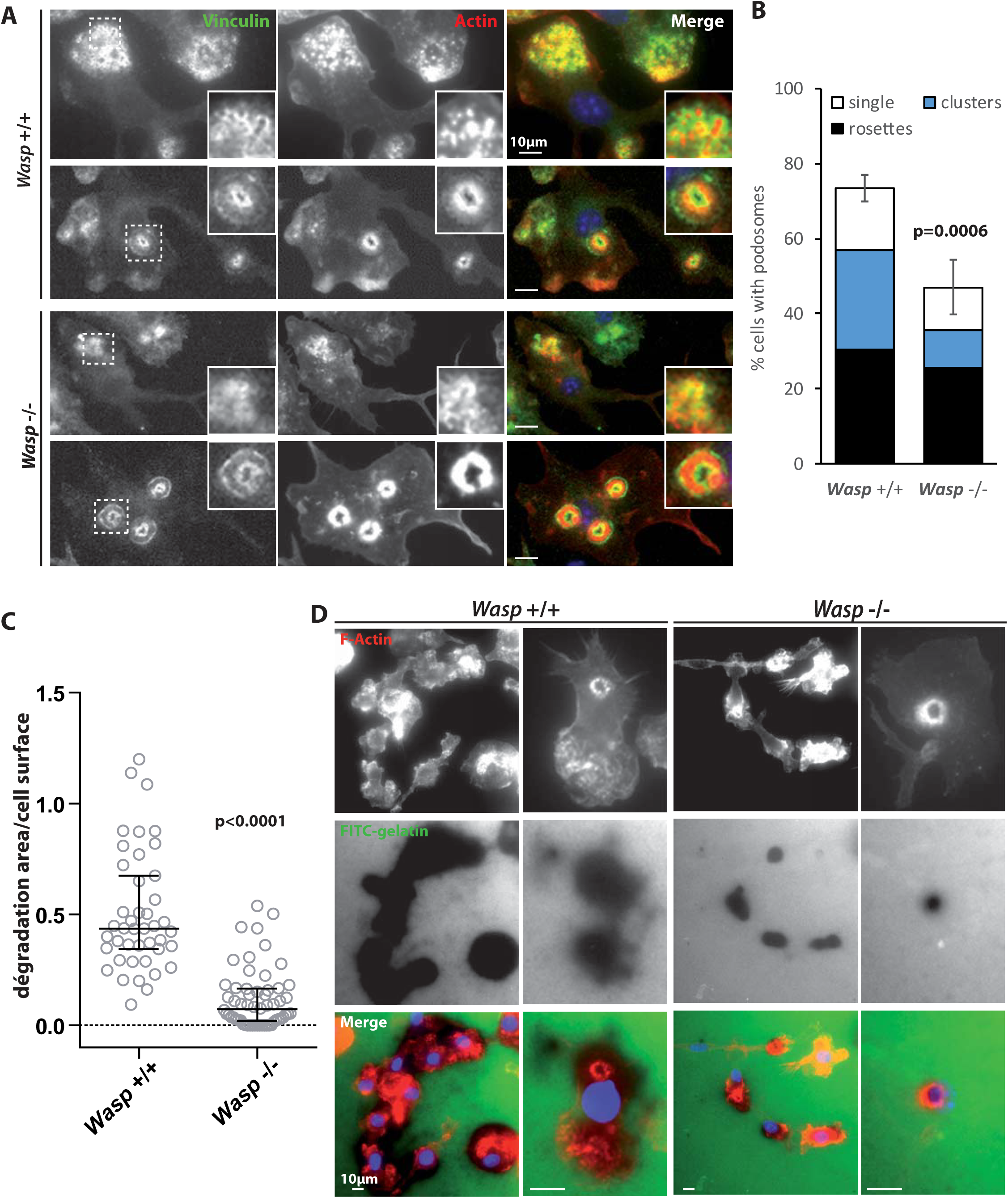
WASP-depleted Hoxb8-macrophages have a defect in podosome organization and ECM degradation. Podosome organization **(A,B)** and FITC-gelatin degradation **(C,D)** were analyzed by fluorescent microscopy and quantified as described in figure 4. **(A, D)** Representative pictures of three independent experiments are shown. **(B)** The percentages of cells with single podosomes, podosome clusters and podosome rosettes in each cell type was quantified on at least 75 cells in 3 independent experiments. **(C)** The ratio of degradation area over cell surface were determined and is expressed as median with interquartile range. Statistics: the p-value obtained with an unpaired student *t*-test is shown.

**Figure 8:**
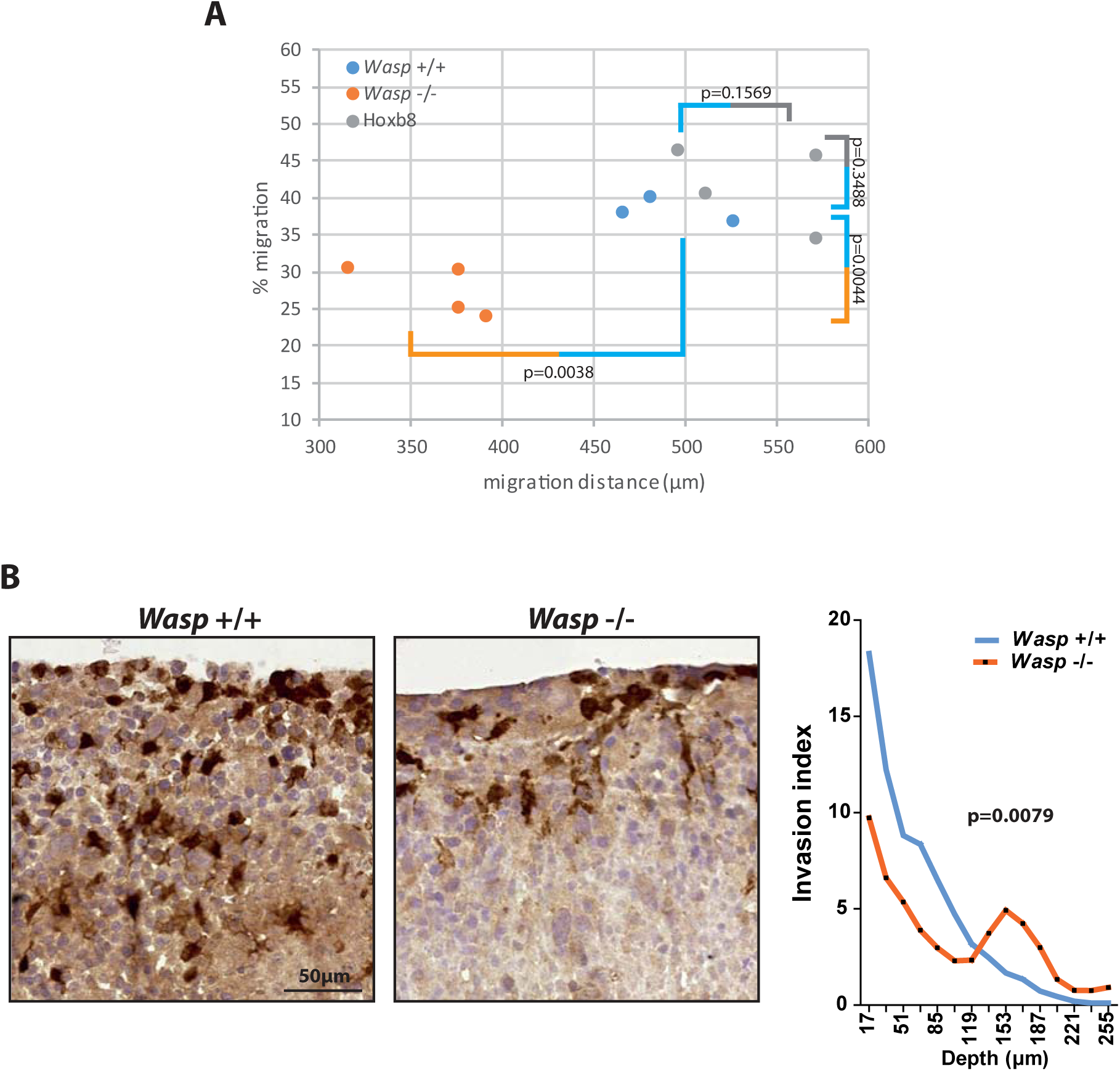
WASP-depleted Hoxb8-macrophages have a defect in 3D migration *in vitro* and *ex vivo*. **(A)** *In vitro* 3D migration of Hoxb8-, *Wasp* +/+ and *Wasp* −/− Hoxb8-macrophages through Matrigel was analyzed as described in Fig. 3. Percentages of cell migration plotted as a function of distance of migration were measured. Mean of duplicates of 3 to 4 independent experiments. Statistics: p-values obtained with paired student *t*-test to compare % of migration (vertical bars) and distances (horizontal bars). **(B)** *Ex vivo* infiltration of cells in LPB-tumors explants was analyzed and quantified as described in Fig. 5. Pictures representative of one out of 3 independent experiments. Statistics: the p-value obtained with an unpaired student *t*-test is shown.

Finally, we tested the capacity of *Wasp* +/+ and *Wasp* −/− Hoxb8-macrophages to infiltrate tumors *ex vivo*. Both the number of tumor-infiltrated *Wasp* −/− Hoxb8 macrophages and the distance of infiltration were reduced (Fig. 8B). These results show that genetic invalidation of *Wasp* affected Hoxb8 macrophages migration in tumor as previously described *in vitro* in Matrigel and tumor cell spheroids for BMDM isolated from *Wasp* KO mice (Park et al., 2014).

In conclusion, these results demonstrate that CRISPR/Cas9-mediated genome editing in Hoxb8-progenitors is highly efficient and represent a potent tool to genetically manipulate macrophage migration effectors.

## DISCUSSION

In the present study, we describe the generation of conditionally immortalized hematopoietic progenitors that can be expanded without limitation, kept in culture, cryopreserved, genetically modified, and that fully retain the potential to differentiate into functional macrophages. Using the CRISPR/Cas9 technology, we show that it provides the opportunity to easily manipulate gene expression, a precious tool to study macrophage biology, and particularly macrophage tissue migration. This is very complementary to the recently published study by Roberts et al. which demonstrates the utility of this model in the study of bacterial pathogenesis (Roberts et al)

Understanding the biology of macrophages is crucial for a large scale of therapeutic contexts such as chronic inflammatory and neurodegenerative diseases, and the majority of solid cancers (Allavena et al., 2008; Condeelis and Pollard, 2006; Friedl and Weigelin, 2008; Yiangou et al., 2006). Macrophage-directed immunotherapy is emerging as a mean to combat cancer and inflammatory diseases. Identification of molecular effectors of macrophage biological functions is thus a requisite to either reduce the damaging effect or boost the protective role of macrophages depending on the concerned disease.

The main caveat in studying macrophages is the lack of tools to manipulate gene expression. SiRNA approaches provide only partial inhibition, require specific investigations for each gene to be knocked down and is often inefficient for the extinction of stable proteins. CRISPR-Cas9 strategies are more appropriate as it allows complete deletion of genes and replacement by a mutant if required. However it is a heavy approach, for cells with a short lifespan that are obtained in a limited number either by drawing blood of humans or preparing bone marrow from mice. Immortalizing progenitors isolated from mouse bone marrow and the subsequent manipulation of gene expression by CRISPR/Cas9 is a powerful alternative with a large scale of applications including studies in vitro, ex vivo and in vivo.

Both our work and the recently published manuscript by Roberts et al. (Roberts et al., 2019) demonstrate the use of HoxB8-conditionaly-immortalized progenitors cells as a powerful tool to study macrophage functions and the possibility of genetically manipulating these cells using the CRISPR/Cas9 technology. Here, we validated the model by analyzing the expression of cell surface differentiation markers, polarization capacity, phagocytosis and production of reactive oxygen species, migration, podosome formation and ECM degradation capacities, while the study by Roberts et al. examined complementary properties of HoxB8 macrophages in infectious settings (Roberts et al., 2019).

We report that the phenotypes of BMDMs and Hoxb8 macrophages were similar regarding both differentiation markers and functions including cell adhesion, macrophage polarization, phagocytosis and ROS production. The only minor difference that we observed was a less efficient mesenchymal migration of Hoxb8 macrophages through Matrigel *in vitro* which corroborated with a diminished ability to degrade FITC gelatin. However, this reduced ability did not impact their capacity to infiltrate tumor tissue *ex vivo*, which was slightly increased compared to BMDMs. This discrepancy underlines the differences that can be observed between *in vitro* experiments performed on acellular ECMs and *ex vivo* experiments performed on tissues isolated from living animals.

In our hands, Hoxb8 macrophages can be kept in culture for three to four months until their migration capacity and macrophage differentiation potential deteriorate. They can be frozen away which allows one to work with the same source of cells and reduces the number of sacrificed mice. This restricted time is in agreement with the 16 weeks delay described by Hammerschmidt and coworkers (Hammerschmidt et al., 2018) and with the standard time that cell lines can be kept in culture without alteration of their behavior. It provides sufficient time for multiple genetic manipulation but limits the possibility to select sub-clones of the different genetically modified cell lines. So, developing advanced culture procedures in the future to optimize maintenance of Hoxb8 immortalized progenitors and extend the culture delay would greatly improve the cell system proposed herein.

Within the scope of our study, we propose Hoxb8 cells as a source of genetically modifiable macrophages for the identification and investigation of tissue migration effectors. As a proof of concept, we deleted WASP, a known actor of migration that partially disrupts podosomes (Isaac et al., 2010; Linder et al., 1999; Park et al., 2014; Vijayakumar et al., 2015). By combining this cellular tool and our *ex vivo* tissue migration assay, we describe a powerful experimental model that is adapted to large scale screening of molecular effectors involved in macrophage migration. This could allow to define the macrophage migration mode within a large collection of healthy or pathological tissues isolated from different mouse models. The use of Hoxb8 cells injected in blood circulation to study tissue recruitment of leucocytes from blood has been recently described (Gran et al., 2018). A future objective will be to examine by intravital microscopy the migration of macrophages in tumors (Gui et al., 2018) by labelling genetically modified Hoxb8 macrophages and directly injecting them within tumors generated in dorsal window chambers. Using this approach, the infiltration of blood monocyte/macrophages into tumors will be by-passed, including the step of monocyte trans-endothelial migration.

Besides the exploration of gene function by deletion, CRISPR/Cas9-mediated genome editing of Hox-B8-immortalized hematopoietic precursor cells also provides the opportunity to easily perform multiple gene deletion, gene overexpression or expression of specific mutants or reporter genes (Salsman and Dellaire, 2017). Added to the capacity of Hoxb8 progenitors to differentiate into a large scale of hematopoietic cells such as DCs (Hammerschmidt et al., 2018), granulocytes (Wang et al., 2006), osteoclasts (Zach et al., 2015) or even B and T lymphocytes (Redecke et al., 2013), it represents a potent, rapid and simple model that opens unlimited possibilities for analysis of leukocyte biological functions including tissue migration.

## MATERIALS AND METHODS

### Animals

C57BL/6 wild-type mice were bred and housed in the accredited research animal facility of the Institute of Pharmacology and Structural Biology (IPBS) that is fully staffed with trained husbandry, technical, and veterinary personnel. All experiments were performed according to animal protocols approved by the Animal Care and Use committee of the IPBS.

### Generation and cell culture of ER-Hoxb8 Progenitor Cell Lines

ER-Hoxb8 cells were generated as previously described (Wang et al., 2006). Briefly, bone-marrow cells were harvested from C57BL/6 mice and hematopoietic progenitors cells were purified by centrifugation on a cushion of Ficoll-Paque. 5×10^5^ cells were pre-stimulated for two days with complete RPMI 1640 medium supplemented with 15% FCS, 1% Penicillin-streptomycin, 1% Glutamin, mouse IL-3, mouse IL-6 and mouse SCF (10 ng/mL each) (Peprotech) in a 6-well culture plate. 2.5×10^5^ cells were seeded on fibronectin coated 12-well culture plate in Myeloid medium (complete RPMI 1640 medium, mouse-GM-CSF (20 ng/mL), β-estradiol (1 µM)) and transduced with 1mL ER-Hoxb8 Retrovirus by spinoculation (1000 *g*, 90 min, 22°C) in the presence of polybrene (24 µg/mL). Polybrene was diluted serially by exchanging half-medium to obtain a final concentration of 1.4 µg/mL over several days. Antibiotic selection of transduced ER-Hoxb8 cells was performed 3 days post infection with G418 (1 mg/mL) and maintained at least two weeks. Immortalized ER-Hoxb8 cells were maintained and enriched in Myeloid Medium (supplemented or not with G418) by serial passages of non-adherent cells every three to four days.

### Differentiation of immortalized ER-Hoxb8 cells toward macrophages

For macrophage differentiation, immortalized ER-Hoxb8 cells were harvested, washed twice with PBS to remove GM-CSF and β-estradiol and cultured at 1×10^6^ cells in complete RPMI 1640 medium supplemented with m-M-CSF (20ng/mL) on glass coverslips in P6-well culture plate for 7 days.

### Vector Construction and Viral Particle Production

gRNA sequence against *Wasp* gene were designed using Benchling.com web tool for genome editing, and selected with high “on-target” and low “off-target” activity. gRNA sequence targeting exon 1 of *Wasp* gene was obtained (forward: 5’-GCTGAACGGCTGGTCCCCCT-3’; reverse: 5’-AGGGGGACCAGCCGTTCAGC-3’) and was cloned into the LentiCRISPRv2 plasmid using BsmB1 overhangs after hybridization and phosphorylation of the gRNA. After transformation in DH5α-competent bacteria, insertion of the gRNA was confirmed by gel electrophoresis of PCR products and the plasmid was purified using the QiaQuick Gel Extraction kit (Qiagen) following provider’s instructions. For virus production, 4×10^6^ HEK293T cells were seeded in 75 cm^2^ flasks in DMEM supplemented with 10% FCS, 1% Penicillin-streptomycin and 1% Glutamine. The next day, cells were transfected with the lentiCRISPRv2 transfer plasmid (5 µg) and both packaging plasmids (pSPAX2 (3.75 µg) and pMD2G (1.25 µg) with transfection reagent PEI. After 24 h, the medium was replaced and 48 h post-transfection, virus-containing supernatants were harvested, filtered, and used for transduction of immortalized ER-Hoxb8 cells. A lentiCRISPRv2 plasmid without any gRNA was used to generate “empty” viruses as a control.

### Generation of ER-Hoxb8 *Wasp*−/− cells with CRISPR/Cas9

To generate the ER-Hoxb8 *Wasp* −/− cells, 1×10^6^ ER-Hoxb8 cells were seeded on fibronectin coated 6-well culture plates in myeloid medium and transduced with 2mL of viral suspension (described above) by spinoculation (1000 *g*, 90 min, 22°C) in the presence of polybrene (8 µg/mL). Polybrene was serially diluted by renewing half-medium over several days to obtain a final concentration of 1.4 µg/mL. Antibiotic selection of transduced ER-Hoxb8 cells was performed 3 days post-infection with puromycine (10 µg/mL) and maintained at least 2 weeks. Immortalized ER-Hoxb8 *Wasp* −/− cells were maintained and enriched in myeloid Medium with puromycine (2.5 µg/mL) by serial passages of non-adherent cells every 3-4 days into new 6-well culture plates. The same protocol was used to generate *Wasp* +/+ control ER-Hoxb8 cells, except that “empty” viruses were used.

### Preparation and differentiation of bone marrow-derived macrophages

Bone marrow cells were isolated from femurs and tibias of C57BL/6 wild-type mice, cultured and differentiated as described (Cougoule et al., 2010).

### SDS-PAGE and immunoblot analysis

Equal amounts of total cell lysates in Laemmli buffer were separated by SDS-PAGE, transferred to nitrocellulose membranes that were probed with mouse monoclonal anti-vinculin (V9131, Sigma, 1/500), polyclonal rabbit anti-WASP (H-250, sc-8353 Santa Cruz 1/200) and anti-N-WASP (H-100, sc-20770 Santa Cruz, 1/100) antibodies and revealed by an enhanced chemiluminescence system (Immobilon TM Western, Millipore,U.S.A.).

### Scanning electron microscopy

ER-Hoxb8 cells and BMDM were cultured on glass coverslips for 16 h, fixed using 0.1 M sodium cacodylate buffer supplemented with 2.5% (v/v) glutaraldehyde and prepared as previously described (Lizarraga et al., 2009) for observation with a JEOL JSM-6700F scanning electron microscope.

### *In vitro* 2D and 3D migration assays

Migration assays were performed in 24-transwells (8-µm pores). Empty transwells were used for 2D migration (Cougoule et al., 2012). Transwells were loaded with either 100 µl of Matrigel (Corning) or 120 µl Fibrillar Collagen (Collagen 2.15 mg/mL (Nutragen), MEM 10x, H_2_O, Bicarbonate buffer) and used for 3D migration (Van Goethem et al., 2010). Matrices were allowed to polymerize for 30 minutes at 37°C, and rehydrated for 3 hours with RPMI 1640 supplemented with 1% Penicillin-streptomycin and 1% Glutamine. Lower chamber were filled with RPMI 1640 containing 10% FCS and 20 ng/mL mMCSF, and upper chamber with RPMI 1640 containing 1% FCS and 20 ng/mL mouse-MCSF. For migration inhibition assay, medium of upper and lower chambers were supplemented with Y27632 20 µM (VWR) or a cocktail of protease inhibitors: 6 µM Leupeptine (Sigma), 0.044 TUI/mL Aprotinine (Sigma), 2 µM Pepstatine A (Sigma), 5 µM GM-6001 (VWR) and 100 µM E64C (Peptides international). Vehicle was loaded as a control. Macrophages (6×10^4^) were serum starved for 3 hours and seeded in the upper chamber. For 2D migration, after 16 h of migration, cells that did not migrate through the porous membrane were removed and cells that migrated underneath the membrane were fixed, stained with DAPI and counted as previously described (Cougoule et al., 2012). For 3D migration assays, after 48 hours of migration, z-series of images were acquired at the surface of the matrices (Top in Fig.3) and inside the matrices with 30 µm intervals (Inside in Fig.3). Experiments were performed in duplicate. Acquisition and quantification of cell migration was performed using the motorized stage of an inverted video microscope (Leica DMIRB, Leica Microsystems, Deerfield, IL), the Metamorph software and the cell counter plugin of the ImageJ software as described (Van Goethem et al., 2010). Migration distance was the maximal migration depth crossed by macrophages inside the matrix. Percentage of migration was obtained as the ratio of the number of cells within the matrix to the total number of counted cells as described (Van Goethem et al., 2010). For live cell imaging of 3D migration (see movie 1 to 3 in Supplementary Material), pictures at the matrix surface and at 300 μm below the surface were recorded every 10 min during 20 h, using the 10× objective of an inverted video microscope (Leica DMIRB, Leica Microsystems, Deerfield, IL, USA) equipped with an incubator chamber to maintain constant temperature and 5% CO2 levels.

### Immunofluorescence microscopy

6×10^4^ macrophages were seeded on coverslips for 24 h. Cells were fixed with 3.7 % paraformaldehyde (Sigma), permeabilized with 0.1% Triton X-100 (Sigma) and stained with anti-vinculin antibody (Clone V9131, dilution 1/100, Sigma) followed by Alexa Fluor 488 anti-mouse Immunoglobulin G Fab2 (dilution 1/500, Cell Signaling Technology) and Texas Red-coupled phalloidin (Dilution 1/200, Molecular Probes Invitrogen). Slides were visualized with a Leica DMLB fluorescence microscope as previously described (Van Goethem et al., 2011).

### FITC-Gelatin degradation assay

Coverslips were coated with 0.5 µg/mL poly-L-lysine and 0.2 mg/mL FITC-gelatin (Invitrogen). Macrophages (6×10^4^) were cultured 24 h on FITC-gelatin, fixed and processed for F-actin staining. Quantification of matrix degradation was performed as previously described (Cougoule et al., 2010).

### Polarization towards M1 or M2 macrophages

ER-Hoxb8 macrophages (5×10^5^cell/well in P6-well plate) were incubated in complete RPMI 1640 medium supplemented with LPS (1 µg/mL) and γ-IFN (20 ng/mL), or IL-4 (20 ng/mL), for macrophage polarization towards M1 or M2 macrophages, respectively as previously described (Cougoule et al., 2012). After overnight culture, cells were harvested and analyzed by FACS as described (Gui et al., 2018).

### FACS analysis

Macrophages were incubated with the following fluorochrome-conjugated monoclonal murine antibodies: anti-Ly6C-FITC (AL-21, BD Pharmingen), anti-CD11b-PE (M1/70.15, Immunotools), anti-F4/80-APC-Cy7 (BM8, Biolegend), anti-CD115-APC (CSF-1R)(AFS98, Biolegend), anti-CD206-PE-Cy7 (MR) (C068C2, Biolegend), anti-CD86-BV650 (GL-1, Biolegend), anti-CD80-PE-Dazzle (16-10A1, Biolegend), anti-Nos2-PE (C-11, Santa Cruz). The acquisition was done with LSR Fortessa (Becton Dickinson) flow cytometer under BD FACS Diva software. For every analysis, 1-2 × 10^4^ cells were acquired. Data were analyzed with FlowJo software (Tree Star) as described (Gui et al., 2018).

### ROS production assay

To monitor the intracellular accumulation of ROS by microscopy, the general oxidative stress indicator, CM-H2DCFDA, the carboxylated analog of the cell-permeant agent 2’,7’-dichlorodihydrofluorescein diacetate (H2DCFDA) (Thermofisher) was used. BMDM and ER-Hoxb8 cells (1.5×10^5^ cells/well on 24-well plate) were primed with LPS 100ng/mL overnight in complete RPMI medium. Cells were washed with phenol-free RPMI and incubated with 200 µg/mL zymosan for 30 min at 37°C. Cells were fixed and slides were visualized with a Leica DM-RB fluorescence microscope.

To quantitatively evaluate the NADPH oxidase activity, the superoxide dismutase (SOD)-inhibitable cytochrome C reduction assay was performed as previously described (Le Cabec and Maridonneau-Parini, 1995). BMDM and ER-Hoxb8 cells (5×10^5^ cells/well on 24-well plate) were primed with LPS 100 ng/mL overnight in complete RPMI medium. Supernatant were removed and cells were incubated with cytochrome C (15 mg/mL) with or without SOD (10000 U/mL) and zymosan (500 µg/mL) in phenol-free RPMI medium for 90 min at 37°C. Reaction were stopped on ice and the reduction rate of oxidized cytochrome C was monitored spectrophotometrically at 550 nm.

### Phagocytosis assay

5 × 10^4^ cells were plated on glass coverslips in RPMI medium at 37°C overnight. Next day, the medium was washed away and FITC-labeled zymosan particles were added to cells at the MOI of 2, in serum-free medium. After a 5-hour incubation, cells were washed in 5% BSA (Euromedex)-containing PBS, then fixed with 3.7 % paraformaldehyde (Sigma), permeabilized with 0.1% Triton X-100 (Sigma), and stained for F-actin with Texas Red-coupled phalloidin (1/200, Molecular Probes) and for nuclei with DAPI. Slides were visualized with a Leica DMLB fluorescence microscope as described (Le Cabec et al., 2002).

### Tumor induction for *ex vivo* migration experiments

To generate fibrosarcomas, 10^6^ LPB cells were injected subcutaneously into the mouse flank as described (Gui et al., 2018). The LPB cell line is a highly tumorigenic murine clonal derivative of TBL.CI2, a methylcholanthrene-induced C57BL/6 mouse sarcoma cell line. Tumors were allowed to grow until approximately 1 cm^3^, then mice were sacrificed, and tumors were resected. Tumor samples were embedded in 3% agarose prepared in PBS (Life technology). Slices measuring 500 µm were obtained with a microtome dedicated to live tissues (Price et al., 1998), with the Krumdieck tissue slicer (TSE Systems) filled with ice-cold PBS (Life Technologies) set to medium blade and arm speeds. Slices were cultured on a 30-mm cell culture insert featuring a hydrophilic PTFE membrane (0.4 µm pore size, Merk Millipore) placed inside 6-well plates containing 1.1 mL complete RPMI 1640 medium with or without 10 µmol/L BB-94 or 20 µmol/L Y-27632. A 5-mm diameter stainless-steel washer was then placed on top of each tissue slice to create a well for macrophages seeding (Gui et al., 2018). The same day, co-cultures were performed by seeding 5×10^5^ murine macrophages (BMDM or ER-Hoxb8 or *Wasp* +/+ or *Wasp* −/−) pretreated or not for 16 h with DMSO, BB-94, or Y-27632 on top of tumor slices and incubated in 37°C, 5% CO_2_ environment for three days. Murine macrophages were labeled with cell tracker (C7025-Green-CMFDA, Invitrogen) according to the manufacturer’s instructions prior to seeding on tissue slices. After 16 h of co-culture, the washer was removed. RPMI 1640 complemented with indicated inhibitors was replaced daily for 3 days before overnight fixation with formalin at 4°C (Sigma-Aldrich).

### Intravital microscopy

Dorsal window chamber surgery and tumor induction for intravital experiments was performed as described (Gui et al., 2018). Before attachment of the coverslip onto the window frame, 20 µL of DMEM containing 1×10^6^ LPB-GFP cells (Life Technologies) and 1×10^6^ Hoxb8 macrophages labeled with cell tracker red CMPTX (Molecular probes) were co-injected subcutaneously in the remaining connective tissue of the lower skin layer. After three days, cells were imaged. Intravital microscopy was carried out on a customized stage for holding mice using a 7MP upright multiphoton microscope (Zeiss) equipped with a 20x/1.0 objective (df=1.8 mm) and a Chameleon-Ultra II laser (Coherent). Animals were anaesthetized by isoflurane inhalation throughout the imaging session. Animal temperature was maintained at 37°C with an Air-Therm-heated environmental chamber and a heating blanket placed under the mouse. The tumor, localized by the fluorescence of LPB cells expressing GFP, was positioned under the objective and cell tracker-labeled Hoxb8 macrophages as well as collagen by second-harmonic generation were observed using a 565-610 nm BP filter. 3D-image stack of sections with z-spacing of 2 µm was acquired every 3 min for two hours to assess 3D cell motility in dynamic z-stack time series.

### Immuno-histo-chemistry on tissue slices

Tissue slices used in co-cultures with macrophages were embedded in paraffin and processed as previously described (Gui et al., 2018). Sections were stained with anti-fluorescein (ab6556 Abcam) and a peroxidase-coupled secondary antibody. Macrophage infiltration was quantified as previously described (Gui et al., 2018).

### Statistical Analysis

Statistical differences were analyzed with the two-tailed paired or unpaired Student’s t-test using GraphPad Prism 6.0 (GraphPad Software Inc.). P-values are indicated in each figure. P<0.05 was considered statistically significant.

## Aknowledgements

We gratefully acknowledge David B. Sykes for the generous gift of Hoxb8 tools, and Etienne Meunier for the generous gift of CRISPR/Cas9 tools and their advices. We acknowledge Yoann Rombouts for his expertise in macrophage polarization. We acknowledge the Toulouse Reseau Imagerie platform, the Bivic facility and Anexplo facility. We acknowledge Isabelle Fourquaux from the CMEAB (Faculté de médecine de Rangueil, Toulouse, France) for SEM experiments.

