## supplemental legend of fig S1 and movies for "Genetic engineering of hoxb8 immortalized hematopoietic progenitors: a potent tool to study macrophage tissue migration"

### Supplemental data

**Supplemental Figure S1:**

**(A)** Proliferation of Hoxb8 progenitors was evaluated by quantifying cell number at day 0 and day 3. **(B)** Morphology of untreated Hoxb8-macrophages (M0) or Hoxb8-macrophages polarized toward the M1 or M2 status were observed by phase-contrast microscopy. Images representative of three independent experiments. **(C)** Migration of BMDMs and Hoxb8 macrophages in 2D was analyzed. The 2D migration index was calculated as the number of nuclei that crossed the polycarbonate membrane of the transwell. Quantification of three independent experiments performed in duplicate (median with interquartile range) is shown. **(D)** Expression level of N-WASP in original Hoxb8-, *Wasp* +/+ and *Wasp* -/- Hoxb8-macrophages was analyzed by Western blot.

**Movie 1: BMDM and Hoxb8 macrophages migrating through fibrillary collagen.** BMDMs or Hoxb8 macrophages were seeded in the upper chamber of transwell chambers filled with fibrillar collagen I. Z-series of images were acquired 1h after seeding at the surface of the matrices, at the surface and at 300 µm below the surface of the collagen gel. Time-lapse imaging was acquired every 10 min during 20 h using the 10× objective of an inverted microscope. Left panels show macrophages at the surface (z=0µm) of the gel over time. Right panels show macrophages in the thickness of the matrix (z=-300µm). Note that cells are progressively disappearing from the upper layer of the gel and concomitantly appearing inside the matrix.

**Movie 2: BMDM and Hoxb8 macrophages migrating through Matrigel.** The same experience as described for Movie 1 with Matrigel instead of fibrillary collagen. Time-lapse every 10 min during 20 h at different z-depth, using the 10× objective of an inverted video microscope are shown.

**Supplemental movie 3: BMDM and Hoxb8 macrophages migrating through fibrillary collagen and Matrigel.** In the same experiments than those described in movie 1, single cells were followed during their displacement inside the matrices. Time-lapse every 10 min during 20 h at different z-depth, using the 10× objective of an inverted video microscope are shown.

**Movie 4: Hoxb8 macrophages migrating in an LPB tumor *in vivo*.** Images were acquired by two-photon microscopy every 3 min during 2 h. Dynamics of Hoxb8 macrophages (labeled with cell tracker, shown as white on the left panel and red on the right panel) together with collagen fibers (imaged by second harmonics, represented in cyan) are shown.
