## Supplementary figures and images for "Genetic engineering of hoxb8 immortalized hematopoietic progenitors: a potent tool to study macrophage tissue migration"

### supplemental Fig S1

#### Supplemental figure S1

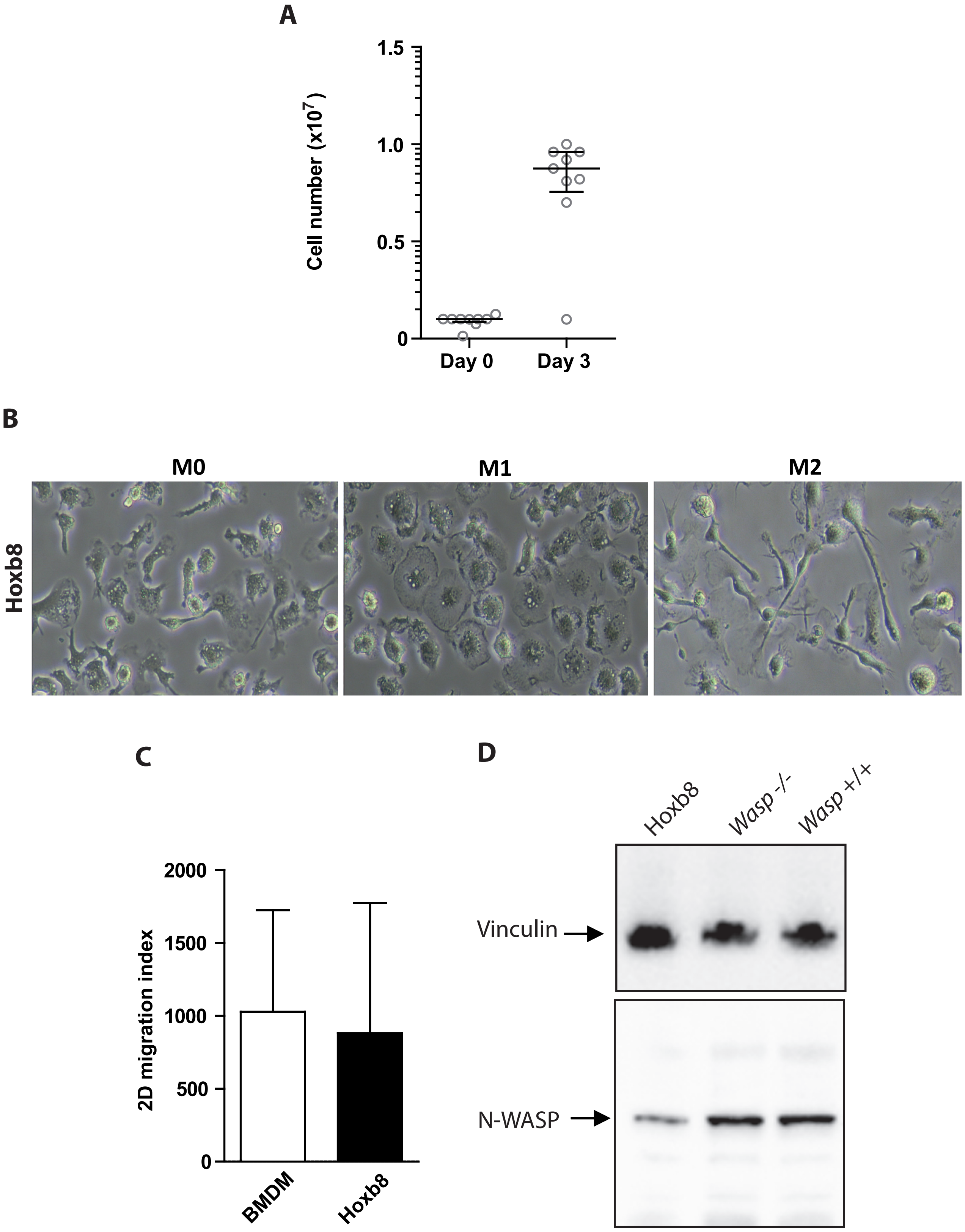
